# Magnetogenetic closed-loop reduction of seizure activity in a rat model of epilepsy

**DOI:** 10.1101/2022.08.19.504501

**Authors:** Abigael C. Metto, Assaf A. Gilad, Galit Pelled

## Abstract

On-demand neurostimulation has shown success in epilepsy patients with pharmacoresistant seizures. Seizures produce magnetic fields that can be recorded using magnetoencephalography. We developed a new closed-loop approach to control seizure activity based on magnetogenetics using the electromagnetic perceptive gene (EPG) that encodes a protein that responds to magnetic fields. The EPG transgene was expressed in inhibitory interneurons under hDlx promoter and kainic acid was used to induce acute seizures. In vivo electrophysiological signals were recorded. We found that hDlx EPG rats exhibited a significant delay in the onset of first seizure (1142.72 ± 186.35s) compared to controls (644.03 ± 15.06s) and significantly less seizures (4.11 ± 1.03) compared to controls (8.33 ± 1.58). These preliminary findings suggest that on-demand activation of EPG expressed in inhibitory interneurons suppress seizure activity, and magnetogenetics via EPG may be an effective strategy to alleviate seizure severity in a minimally invasive, closed-loop and cell-specific fashion.

## Introduction

Mesial temporal lobe epilepsy (TLE) is the most common type of epilepsy with over 28,000 new diagnoses in the United States every year ^1^. TLE remains difficult to manage in 33-44% ^1-3^ of patients causing a significant increase in their mortality rates by 4-7 times that of the general population ^4^. TLE is characterized by focal-onset seizures ^5^ with the majority of them originating in the hippocampus ^6,7^. The first-line treatment for symptomatic epilepsy is use of antiseizure medications (ASMs) which facilitate inhibition or diminish excitation in specific targets responsible for seizure generation. The mechanism that these drugs work is through actions at the receptor sites on target neuronal membranes to regulate neurotransmitters, ion channels, membrane permeability to various ions ^3,8^, neuromodulator function ^3^, and intracellular signaling pathways ^9^. The most common treatment for TLE is carbamazepine or levetiracetam ^3^. Pharmacotherapy administered immediately following the occurrence of the first seizure has been shown to be beneficial in the short-term ^10^. However, in the long-term (> 3 years), there is a greater possibility of seizure recurrence and adverse side effects ^11^. Imperatively, 50-70% of patients with TLE are pharmacoresistant ^12^ and at least a third of this population suffer from inadequately controlled seizures accompanied by disabling psychosocial disturbances ^3,13-16^, which substantially affect their life quality and life expectancy ^17^.

Deep brain stimulation (DBS) has emerged as an alternative therapy for pharmacoresistant epilepsy patients who are not candidates for resective surgery. DBS delivers electrical stimulation to target brain areas via implantable electrodes ^18^ and is commonly used to mitigate symptoms of Parkinson’s disease (PD), movement disorders ^19^ and chronic pain ^20^. High frequency stimulation (100-180 Hz) delivered to the subthalamic nucleus (STN) or the hippocampus ^21^ of TLE patients has been successful in a subset of patients who became seizure-free after 6 days.

Recent preclinical research has also focused on the prospect of optogenetics to control seizures originating in the hippocampus. Kim et al. ^22^ demonstrated that closed-loop video-EEG monitoring combined with on-demand optogenetic stimulation in a mouse model of TLE can effectively decrease seizure duration and improve memory. This work had paved the way to engineering new devices that will allow closed-loop and on-demand stimulation of seizures. New strategies that will allow simplified design and minimal invasiveness could transform the field of seizure management.

A potential method to identify the timing and the location of a seizure in real-time is through the magnetic fields that are produced by the seizure’s hyper-synchronized neuronal activity ^23,24^. These magnetic fields are evident by Magnetoencephalography (MEG) which is a functional neuroimaging method that is based on magnetic fields that are produced by electrical currents in the brain. MEG is extremely useful in localization of the epileptogenic zone ^25^ and is often used in the presurgical assessment of patients with epilepsy. Studies showed that MEG is more accurate in detecting epileptiform activity in the lateral neocortex compared to scalp EEG ^26^.

The magnetic fields induced by the seizures open potential new strategies to monitor seizures in real time and to use this as a biomarker for neuromodulation; The magnetic fields may also be used to induce neural activity via magnetogenetics technologies. Here we tested if the magnetic fields induced during a seizure in the established kainic acid (KA) rat model ^27^ could be used both as the sensor and the activator to suppress seizure activity in an animal model. We capitalized on a recently discovered membrane-associated protein that is sensitive to magnetic fields, the Electromagnetic Perceptive Gene (EPG) ^28-31^; it was demonstrated that remote activation of EPG by electromagnetic fields significantly increases intracellular calcium concentrations, indicative of cellular excitability ^32^. Thus, we tested if EPG expressed in inhibitory hippocampal interneurons will be sensitive to the magnetic fields generated during a seizure, that in turn will activate the inhibitory interneurons which will shut down or disrupt the circuit, interrupting seizure activity and progression in a closed-loop and cell-specific fashion.

## Results

Intrahippocampal injection of KA is an established model to induce acute seizures in rats ^27^. We tested the effectiveness of EPG expression in the hippocampal CA3 neurons to reduce seizure latency, frequency and duration in this model. Adeno-associated virus (AAV9) encoding for EPG (experimental, hDlx EPG, n=9) or green fluorescence protein (GFP) (control, hDlx GFP, n=9) under hDlx promoter which is specific to inhibitory interneurons^33^ was stereotaxically injected into the CA3 region of the hippocampus in the right hemisphere (coordinates according to Bregma: Anterior-Posterior (AP) 4 mm, Medial-Lateral (ML) 3 mm, Dorsal-Ventral (DL) -3.1 mm). Three weeks following virus delivery, rats were anesthetized with isoflurane (0.5%) and dexmedetomidine (1.5mg/ml/hr) and seizures were induced using intrahippocampal injection of 1µl KA (0.5µg/ 0.1 µl) into the right CA3. Figure 1a shows an illustration of the experimental design. Figure 1b shows the putative mechanism of the closed-loop seizure suppression.

**Fig. 1:**
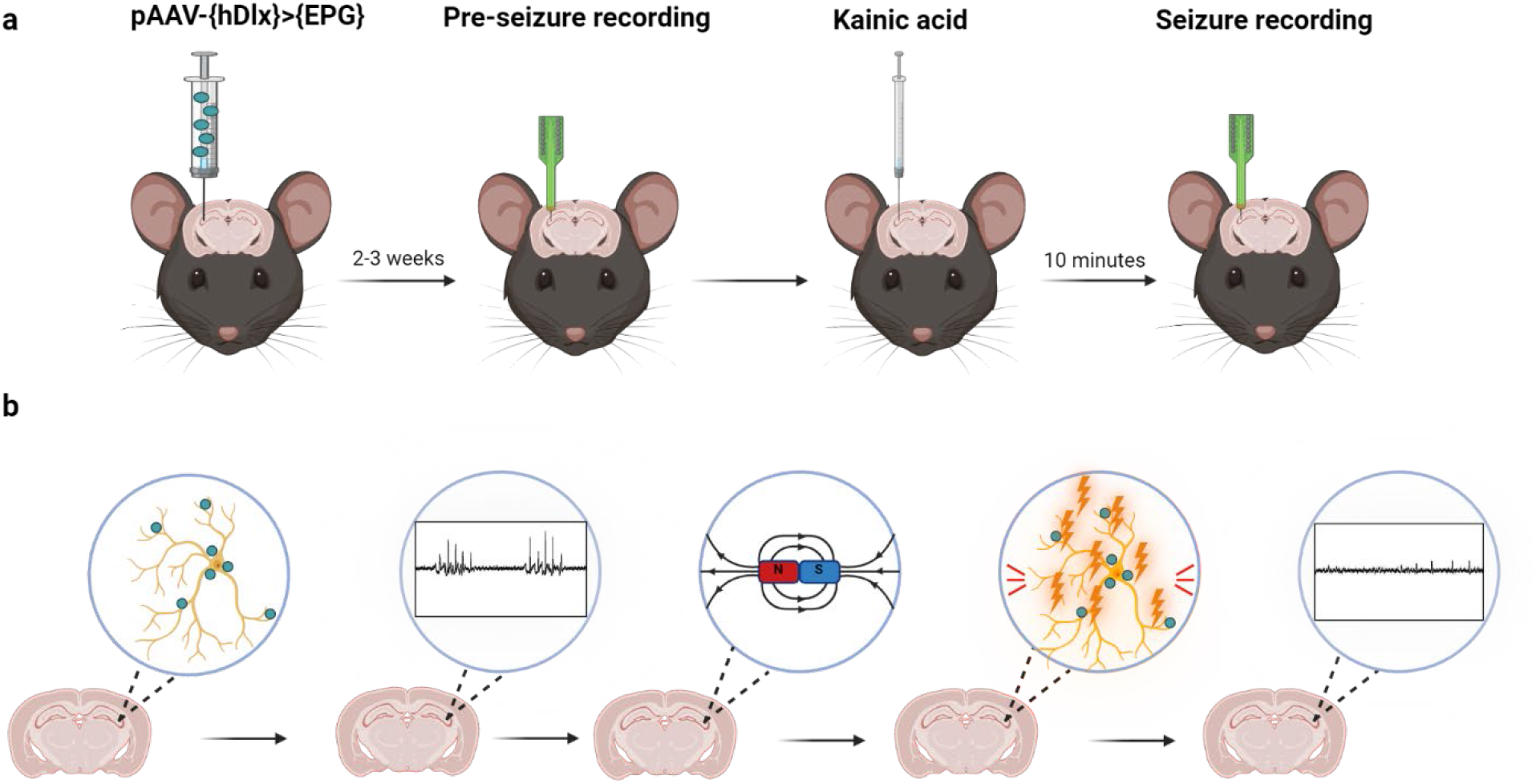
EPG is activated in response to magnetic fields generated by neuronal firing during seizures in the hippocampus. **a**. Schematic illustration of the experimental design: Viral injection (AAV9) of EPG targeting inhibitory neurons in the CA3 region of the right hippocampus in adult rats. EPG is expressed in neurons within 2-3 weeks. Kainic acid injected into the same region was used to induce acute seizures. **b**. Suggested mechanisms of closed-loop suppression of seizure(s): When a seizure occurs, firing of neurons produces electrical currents that generate magnetic fields, which in turn activate EPG, subsequently terminating seizures.

Tungsten electrodes were positioned in the same location and extracellular recordings of local field potentials (LFP) were acquired using Spike2 system, at a sampling rate of 20kHz. Seizures amplitude, frequency and length were measured. Both the experimental, hDlx EPG rats, and the control, hDlx GFP rats, have exhibited similar seizure charectarisctics. Figure 2 shows representative LFP traces and spectrograms (bottom) recorded from an hDlx EPG rat (a) and an hDlx GFP rat (b).

**Fig. 2:**
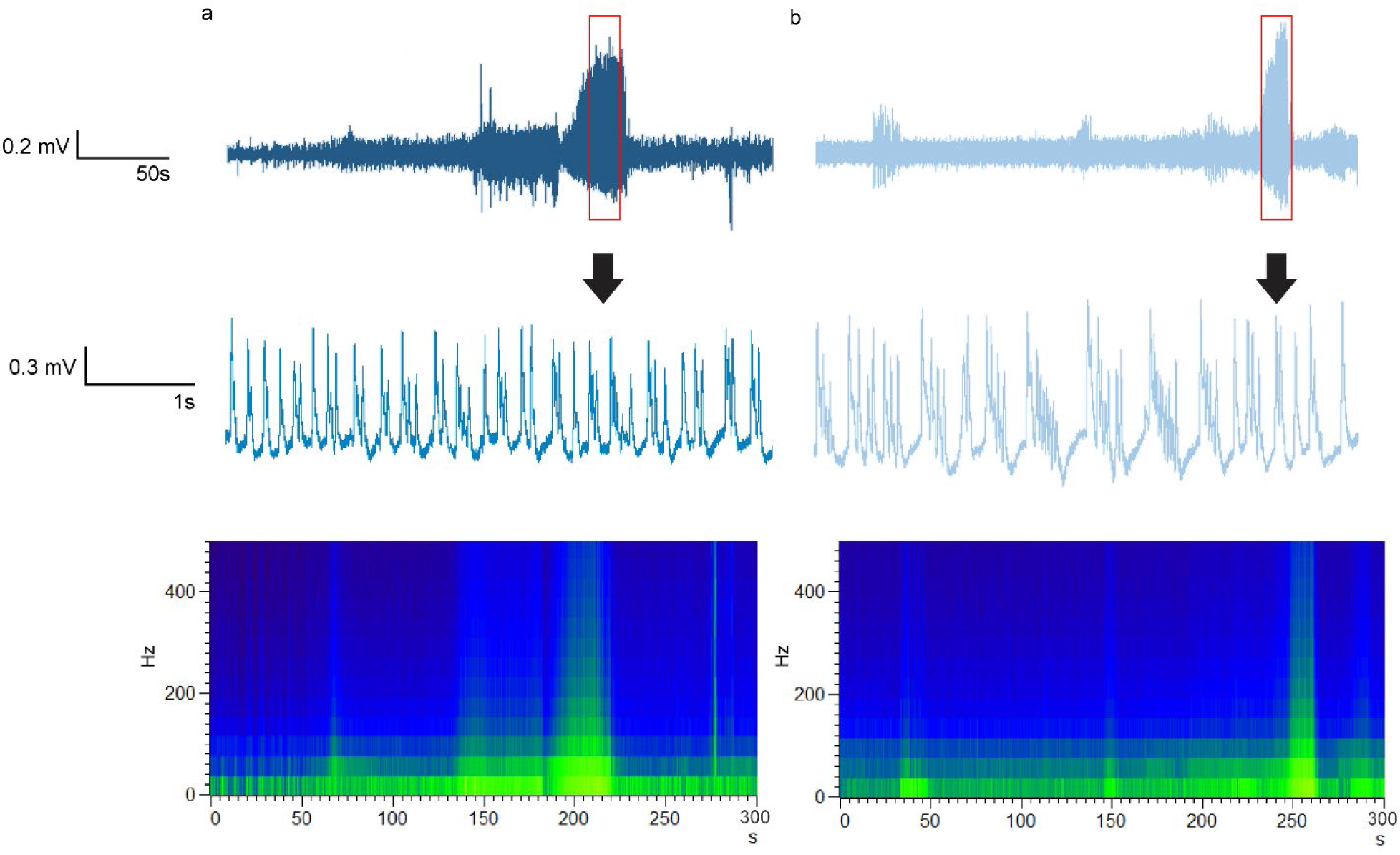
Initiation and progression of seizure over time. **a**. Recordings of extracellular waveforms 10 minutes after injection of KA in an hDlx EPG rat, and in a **b**, hDlx GFP rat. Bottom, Spectrograms representing frequencies of waveforms **a** and **b** respectively, over time.

### EPG in interneurons delays the onset of seizures

Intrahippocampal injections of KA induces acute seizure that appear within minutes after injection ^27^. Figure 3a shows the seizure onset time for the hDlx EPG group and the hDlx GFP group. The onset of seizures was 1142.72 ± 186.35s in the hDlx EPG group and 644.03 ± 15.06s in the hDlx GFP group. D’Agostino & Pearson test for non normal distributions was applied and the data was normally distributed. An unpaired Student t test showed a significance change of P = 0.028.

**Fig 3:**
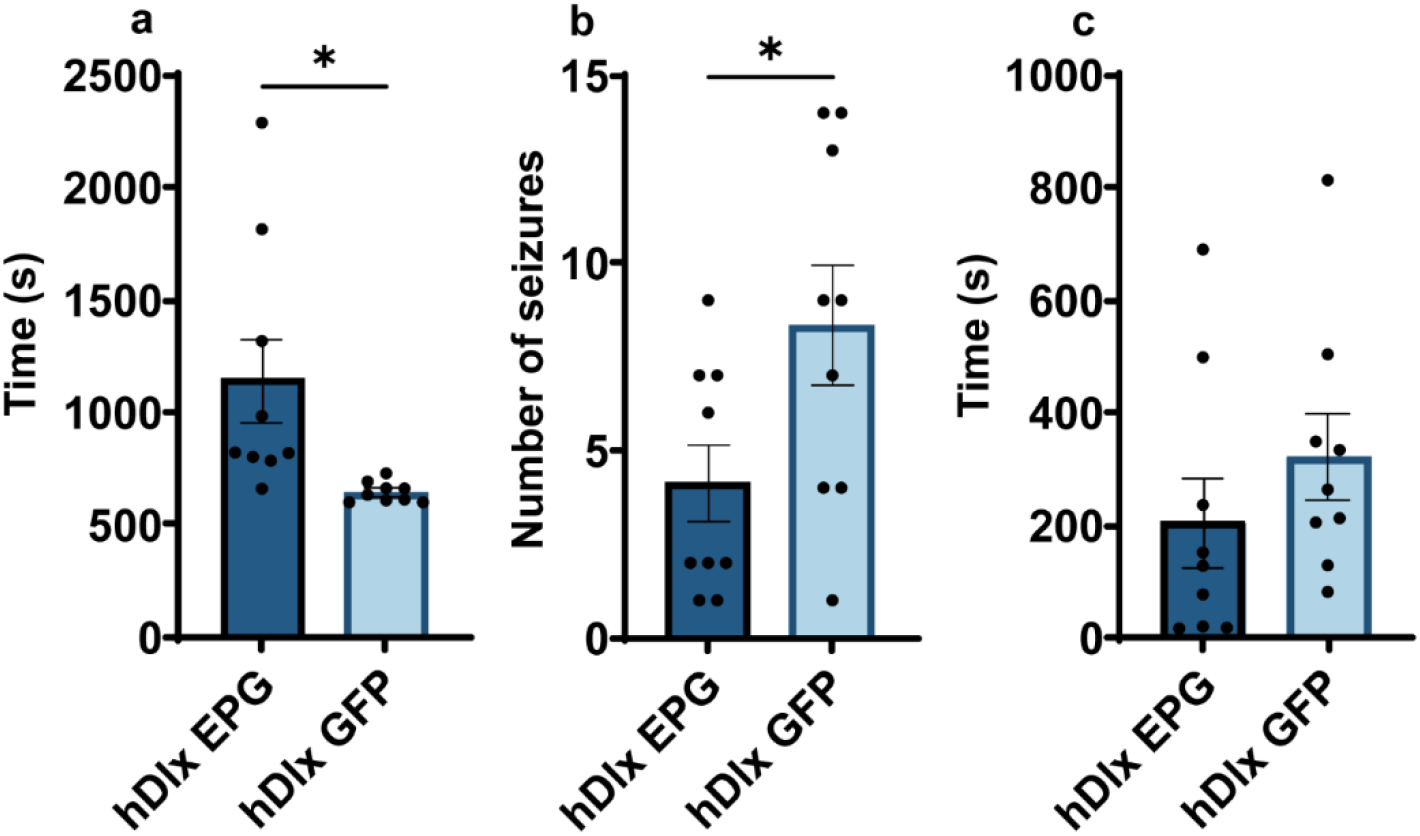
Rats expressing EPG in CA3 inhibitory neurons exhibited significantly longer latency to first seizure and a significantly smaller number of seizures compared to control rats. **a**. Seizure onset: Rats expressing EPG (hDlx EPG, n = 9) in CA3 inhibitory interneurons had a significant delay in the onset of the first seizure with an average time of 1142.72 ± 186.35s compared to control rats (hDlx GFP, n = 9) that had a delay of 644.03 ± 15.06s (*P = 0.028). **b**. Number of seizures; Rats expressing EPG exhibited significantly less seizures (4.11 ± 1.03) compared to control rats (8.33 ± 1.58) (*P = 0.042). **c**. Total seizure duration; Rats expressing EPG had a seizure duration of 203.37 ± 79.22s while control rats had a duration of 320.62 ± 74.43s, P = 0.352.

### EPG in interneurons decreases the number of seizures

Figure 3b shows the number of seizures recorded 60 min after intrahippocampal KA injection. The hDlx EPG group experienced significantly less seizures, 4.11 ± 1.03, compared to the hDlx GFP group, 8.33 ± 1.58. D’Agostino & Pearson test showed normal distribution and an unpaired Student t test showed a significant change of P = 0.042. The duration of the seizures was calculated for each event. The D’Agostino & Pearson test showed that the data was not normalaly distributed. Therefore, the differences between groups were analyzed using the Mann-Whitney U test. Figure 3c shows the total duration of seizures. No significant differenc between the hDlx EPG group (203.368 ± 79.22s) and the hDlx GFP group (320.623 ± 74.43s, P = 0.352) was observed. Previous evidence suggest that the lesion induced by the electrode positioning itself can by itself suppress some of the seizures. Therefore, an additional control group that did not express any transgene, but received a similar intrahippocampal KA injection (naive) was tested. When we compared the hDlx EPG group to the naive group we found that the hDlx EPG group had significantly less total seizure duration (203.368 ± 79.22s) compared to the group with no injection (1008.68 ± 176.70s, **P = 0.006) (Fig.3).

**Fig. 4:**
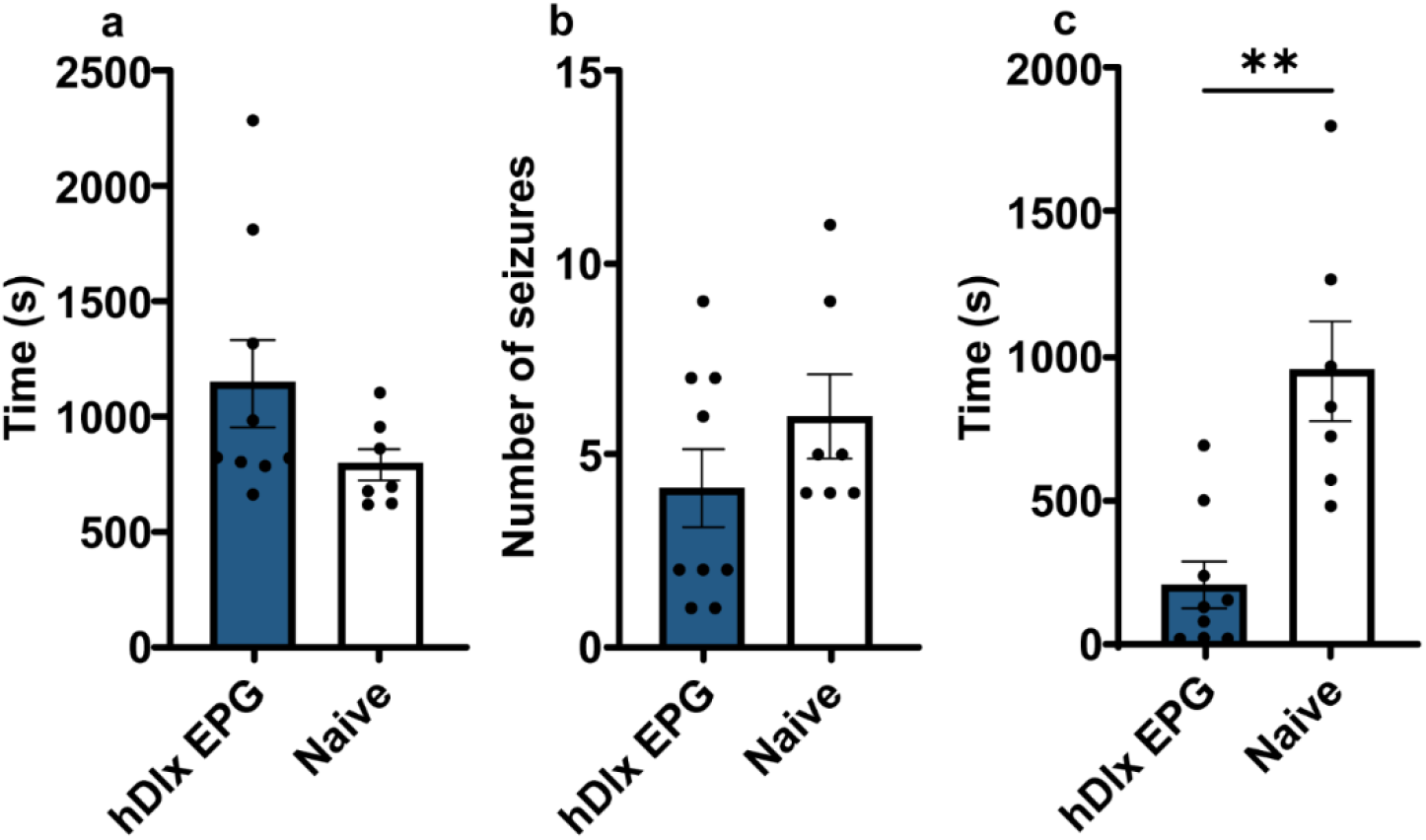
Rats expressing EPG in CA3 inhibitory neurons exhibited significantly shorter total seizure duration compared to naive rats. **a**. Seizure onset: Rats expressing EPG (hDlx EPG, n = 9) in CA3 inhibitory interneurons had an average onset of 1142.72 ± 186.35s compared to naive rats (n = 7) (788.81 ± 70.79s), P = 0.11. **b**. Number of seizures; Rats expressing EPG experienced 4.11 ± 1.03 seizures compared to naive rats (6 ± 1.07), P = 0.23, **c**. Total seizure duration; Rats expressing EPG had a seizure duration of 203.37 ± 79.22s while naive rats had a duration of 1008.68 ± 176.70s, **P = 0.006.

### Immunofluorescence confirmed localization of EPG in inhibitory neurons

Validation of EPG expression in inhibitory inetreneurons was perfomed using immunofluorescence. Following electrophysiology recordeings the rats were perfused and the brain removed. Histology was then performed on 30 µm thick brain slices that were cut in the hippocampus level. Fig. 5 shows immunofluorescence imaging from hDlx EPG rats. GAD67 which is a marker for inhibitory neurons was used to confirm cell specificity, and GFP was used to confirm EPG expression. Microscopy results showed that 88% of neurons that were positive for EPG expression were also positive for GAD67 expression, cofirming EPG expression in interneurons.

**Fig. 5:**
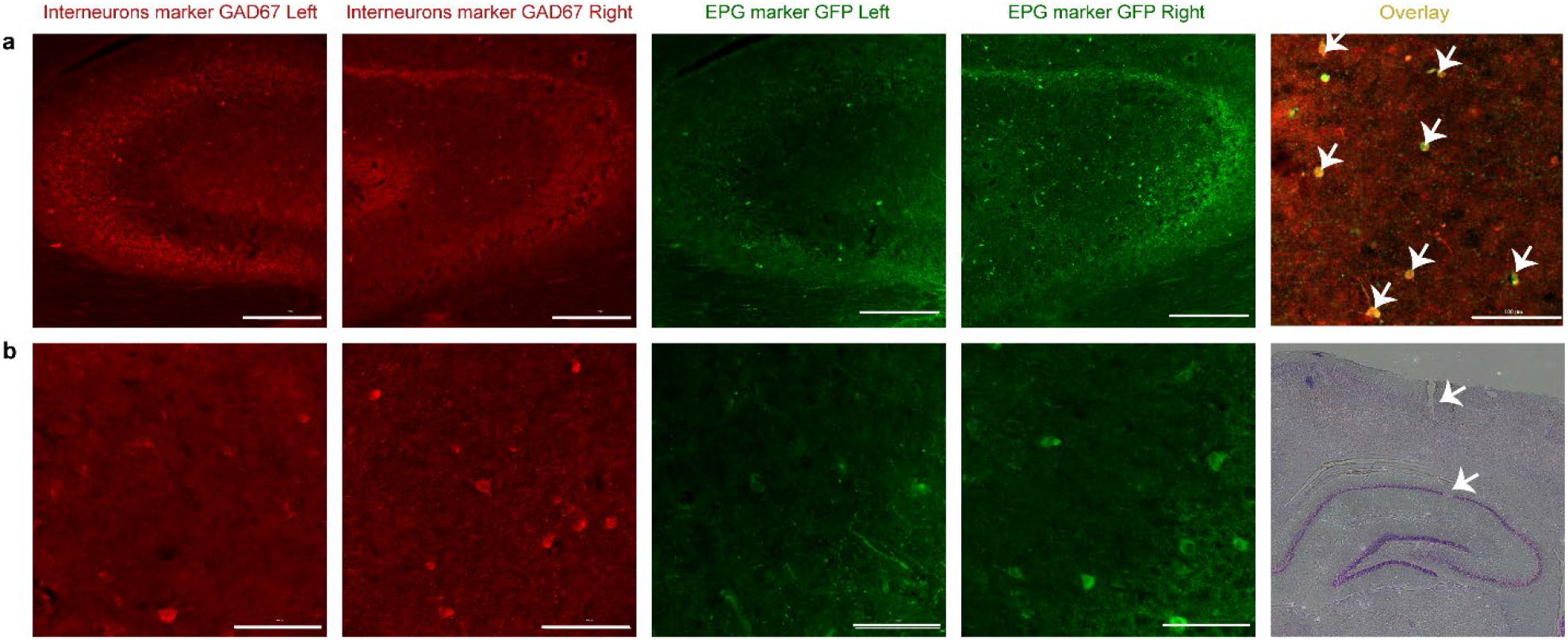
Immunofluorescence showing localization of EPG in CA3 inhibitory neurons. **a**. Inhibitory inetreneurons labled with red and EPG-expressing neurons labled with green in the left and right hippocampus (10X, Scale bar = 300 µm). Far right, overlay of GAD67 and GFP (40X, scale bar = 100 µm). **b**. Higher magnification of inhibitory inetreneurons labled with red and EPG-expressing neurons labled with green in the left and right hippocampus (20X, Scale bar = 200 µm). Far right, Nissl stain showing the recrding electrode tract.

### EPG in excitatory neurons did not affct seizure onset, number and duration

We tested if EPG expression in excitatory neurons will also lead to reduction in seizure severity. For this set of experiments AAV9 encoding for EPG (experimental, CaMKII EPG, n=7) or green fluorescence protein (GFP) (control, CaMKII GFP, n=8) under CamKII promoter which is specific to excitatory neurons was stereotaxically injected into the CA3 region of the hippocampus in the right hemisphere (AP 4 mm, ML 3 mm, DL -3.1 mm). Three weeks following virus delivery, rats were anesthetized with isoflurane (0.5%) and dexmedetomidine (1.5mg/ml/hr) and seizures were induced using intrahippocampal injection of 1µl KA (0.5µg/ 0.1 µl) into the right CA3. No significant differences in the seizure onset, the number fo seizures and seizure duration were found between the CamKII EPG and CaMKII GFP rats.

**Fig. 6:**
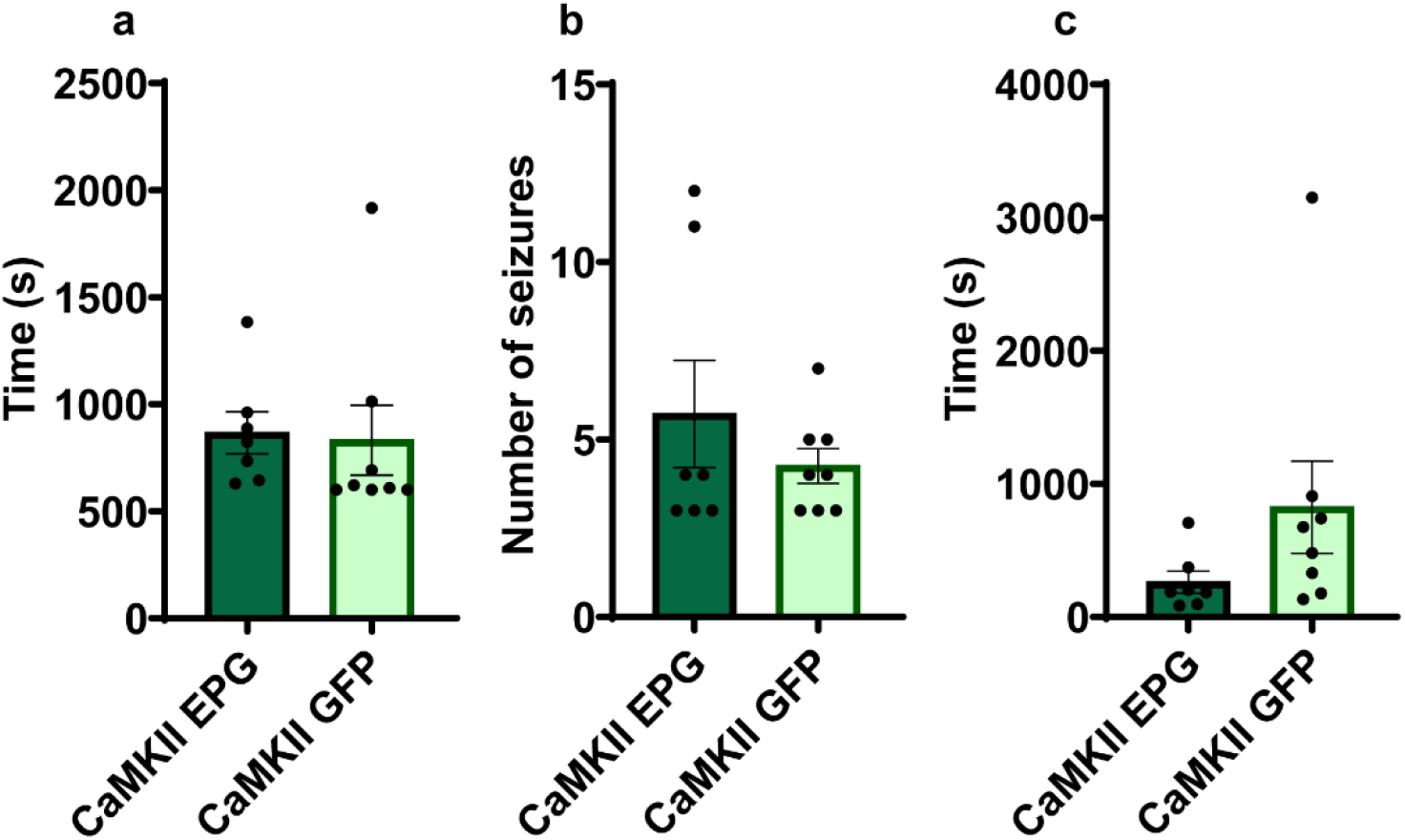
Rats expressing EPG in CA3 excitatory neurons did not exhibited changes in seizure severity. **a**. Seizure onset: Rats expressing EPG in CA3 excitatory neurons showed an average onset of 866.47 ± 97.90s (n=7) compared to control rats who showed an average onset of 831.63 ± 162.80s (n=8), P = 0.15. **b**. Number of seizures; Rats expressing EPG experienced 5.71 ± 1.51 seizures compared to control rats who experianced 4.25 ± 0.49 seizures, P = 0.98, **c**. Total seizure duration; Rats expressing EPG had a seizure duration of 261.81 ± 82.22s while control rats had a duration of 824.14 ± 345.99s, P = 0.12.

## Discussion

In the past decade neurostimulation has been identified as an effective startgy for seizure mangment. Closed loop neurostimlation approaches are necceesay to control seizures and new devices and software that allow real-time seizure detection are being developed. Current hardware that provides closed-loop detection and stimulation requires invasive placement of the apparatus components. New neural engineering approaches combined with molecular biology tools could offer new theraputic startagies to alleviate seizures severity and also provide new tools to understand the fundamental basis and mechanisms of seizures development and seizure therapies.

The premise for the current work rests on the principle that a moving current generates a magnetic field according to Fleming’s right-hand rule. In the epileptic brain, electric currents arise from dendritic action potentials of pyramidal neurons that fire hyper-synchronously and in parallel during epileptiform activity. Using MEG, the source of epileptic spikes and interictal discharges ^34-36^ can be localized in three dimensions and the direction of current flow can be determined ^3^. An advantage of this technique is that magnetic fields are not distorted by the scalp or skull ^37^. Indeed, MEG provides a high temporal and spatial resolution of current sources deep within the brain (for review see ^38^).

Capitalizing on this phenomenon, we investigated the potential for EPG to supress seizure activity in an acute model of TLE. The advantage of this approach is that the EPG will act as both the seizure detection component and the neuronal modulator and will provide a closed-loop system. Previous studies have demonstrated that magnetic stimulation of neurons expressing EPG induced neuronal activity ^28^, and increased calcium influx ^32^. Magnetic stimulation of rats that expressed EPG in their sensorimotor cortex evoked motor activity ^28^ and neuroplasticity ^30^. Here we tested if hippocampal neurostimulation via EPG could effect hippocampal circuitry.

It remains to be determined what are the brain targets for neurostimulation to achive for maximizing therapeutic outcome. In humans, high-frequency stimulation in the range of 100-180Hz targeting the STN ^39^ and hippocampus ^40^ has been shown to be well tolerated and effective in reducing seizure occurrence by 93% and 67%, respectively. Preclinical models to test the effectiveness of DBS in sub-cortical regions have shown similar outcomes ^41^. High frequency stimulation interferes with locally synchronized oscillations ^42^ within the seizure focus and is therefore considered inhibitory. Indeed, clinical evidence suggests that seizure-triggered stimulation, for example, Responsive Neurostimulation (RNS) is an effective strategy ^43^. RNS is based on a closed-loop stimulation paradigm that provides the ability to adjust seizure detection parameters according to each individual patient ^44,45^. Jarociewicz et al. ^45^ found that the most common stimulation parameters in 256 patients consisted of two pulses administered at 100-200 Hz. RNS is advantageous because it results in minimal adverse side effects ^46^.

Furtheremore, targeting different neuroal population could lead to different outcomes. In the hippocampus, 10-15% of neurons are gamma-aminobutyric acid (GABA)ergic inhibitory interneurons and the majority of the other neurons are glutamatergic excitatory pyramidal neurons ^47^. Evidence from preclinical models has shown that increasing hippocampal and STN inhibition with N-methyl-D-aspartate (NMDA) antagonists ^48,49^ and GABA agonists ^50,51^ can suppress seizures by interfering with seizure progression. For example, gene-based therapy designed to enhance GABA resulted in antiseizure effects^48^. Noe et al also demonstrated that AAV-mediated expression of neuropeptide Y, which is an endogenous peptide involved in the modulation of glutamate release from CA3 pyramidal neurons ^51^, had a significant anti seizure effect ^3^. Additionly, in their closed-loop optognetics modulation of seizure activity, Kim et al.^22^ targeted inhibitory ineterneurons in CA3 region. In agreement with previous studies, we found that targeting the inhibitory interneurons leads to an effective way to minimize seizure burden. This approach delayed the onset of seizures and decreased their frequncy. On the other hand, targeting the excitatory neurons did not change seizure charectaristics. This implicates the inhibitory neurons as a good target for seizure management.

Although the number of components for seizure detection and neuronal activation was considerably reduced in this new approach, it still required invasive delivery of the transgene. New and upcoming approaches for minimally invasive gene-delivery that are being designed for a different health disorders, are likely to offer new ways to deliver transgenes that can cross the blood brain barrier and target a specific neural population ^52^.

Finally, it will be important to test if the EPG strategy will be effective not only in an acute seizure model, but also in a chronic seizure disease model, and if targeting STN will also be effective in reducing sizure severity. This new appach can provide new means for adaptive seizure control, and a new tool for understanding seizure charectaristics in animal models.

## Materials and Methods Intrahippocampal AAV Injection

All procedures were approved by the Institutional Animal Care and Use Committee at Michigan State University. Adult Wistar Furth and Long Evans rats (200g-400g, > 10 weeks old, n = 40, (17 = female, 23 = male)) were anesthetized with isoflurane (5% for induction; 2.5% for maintenance and surgery) and positioned in a stereotaxic frame. The microinjection was positioned at AP -4.0 mm, ML 3.0 mm, DV -3.1 mm relative to Bregma ^53^. In the experimental group, Adeno-associated virus encoding for EPG under hDlx promoter (AAV9-hDlx-EPG-IRES-EGFP) was injected into the CA3 region of the right hippocampus. Two control groups were included: A sham group that received an injection of virus encoding only for GFP (AAV9-hDlx-EGFP), and a group that did not receive an injection at all.

### Electrophysiology

Electrophysiology took place two to three weeks after transgene delivery. Rats were anesthetized with isoflurane (5% for induction; 2.5% for surgery; and 0.5% throughout the recording) and a constant infusion of Dexmedetomidine (0.5mg/kg/hr) subcutaneously. Isoflurane was administered at 0-5 % during all recordings to minimize its effect on neuronal activity ^54,55^.

To obtain Local Field Potentials (LFPs), a monopolar tungsten microelectrode (impedance: 200-300kΩ, FHC, Bowdoin, ME) was inserted into the right hippocampus CA3 region. Baseline recording was acquired for ten minutes. Following the baseline recording, kainic acid (KA) (0.5*µ*g/0.1*µ*l) was injected into the right hippocampus and ten minutes after KA injection, recordings were acquired for 1h. The signals were recorded with an Intan Recording System (Intan Technologies, Los Angeles, CA) and Spike2 (Cambridge Electronic Design, Cambridge, UK) at a sampling rate of 20kHz ^56-59^.

### Immunohistochemistry

Rats that developed seizures were perfused transcardially with 1X Phosphate Buffered Saline (PBS) and 4% Paraformaldehyde (PFA). The brains were extracted and placed in 4% PFA overnight after which they were placed in varying concentrations of sucrose cryoprotection solution until they sank (10%, 20% and 30% consecutively). Frozen brains were sectioned at a thickness of 30*µ*m using a Leica CM3050 S Cryostat (Leica Biosystems, Buffalo Grove, IL) and slices were placed in 4C in PBS. Antigen retrieval was achieved by heating the brain slices to 95C for 5 minutes in sodium citrate buffer (10mM sodium citrate, 0.05% Tween 20, pH 6.0). After cooling, slices were washed in PBST (0.1% Tween 20) 3 times and a blocking solution added (donkey serum, Sigma-Aldrich, St. Louis, MO) for 1 hour. Slices were then incubated with the primary antibodies (1:500, Anti-GAD67, ab97739, Abcam, Boston, MA and Anti-GFP, Jackson ImmunoResearch Labs, West Grove, PA) and rocked overnight at 4C. The following day, slices were washed in PBST 3 times after which they were incubated with the secondary antibodies (1:500, Donkey Anti-Rabbit conjugated with Alexa Fluor 647, and Donkey Anti-Chicken conjugated with Alexa Fluor 488, Jackson ImmunoResearch Labs, West Grove, PA), and rocked in the dark for 3h at 4C. Sections were subsequently washed three times with PBS (15min each) and mounting media (with DAPI) was applied. Images were acquired using the Biotek Cytation 5 Cell Imaging Multi-Mode Reader (Biotek Instruments, Inc., Winooski, VT) configured on an inverted microscope.

### Analysis and Statistics

Recordings were analyzed using NeuroScore software (Data Sciences International, St. Paul, MN). Signals were high-pass filtered at 0.1Hz and seizure detection performed with the following parameters: threshold ratio: 1.2, maximal ratio: 20, minimum value: 100*µ*V, minimal spike duration: 1ms, maximal spike duration: 500ms. Seizure events were defined as having at least 4 spikes, minimum interspike interval of 0.05s, maximum interspike interval of 0.6s, minimum train duration of 7.5s, and a train join interval of 1s. Analyses were validated by visual inspection of traces offline. D’Agostino & Pearson test was performed on all data sets to tests for normal distribution of data points. Further statistical tests were selected based on these results.

## Notes

### Competing Interest Statement

The authors have declared no competing interest.

## References

1. Asadi-Pooya, A.A., Stewart, G.R., Abrams, D.J. & Sharan, A. Prevalence and Incidence of Drug-Resistant Mesial Temporal Lobe Epilepsy in the United States. World Neurosurg 99, 662–666 (2017).

2. Asadi-Pooya, A.A., Nei, M., Sharan, A. & Sperling, M.R. Historical Risk Factors Associated with Seizure Outcome After Surgery for Drug-Resistant Mesial Temporal Lobe Epilepsy. World Neurosurg 89, 78–83 (2016).

3. Engel, J. Seizures and epilepsy, (Oxford University Press, Oxford ; New York, 2013).

4. Sperling, M.R. The consequences of uncontrolled epilepsy. CNS Spectr 9, 98-101, 106-109 (2004).

5. Nayak, C.S. & Bandyopadhyay, S. Mesial Temporal Lobe Epilepsy. in StatPearls (Treasure Island (FL), 2022).

6. Tatum, W.O.t. Mesial temporal lobe epilepsy. J Clin Neurophysiol 29, 356–365 (2012).

7. Blumcke, I., Coras, R., Miyata, H. & Ozkara, C. Defining clinico-neuropathological subtypes of mesial temporal lobe epilepsy with hippocampal sclerosis. Brain Pathol 22, 402–411 (2012).

8. Sills, G.J. & Rogawski, M.A. Mechanisms of action of currently used antiseizure drugs. Neuropharmacology 168, 107966 (2020).

9. Landmark, C.J. Targets for antiepileptic drugs in the synapse. Med Sci Monit 13, RA1–7 (2007).

10. Krumholz, A., et al. Evidence-based guideline: Management of an unprovoked first seizure in adults: Report of the Guideline Development Subcommittee of the American Academy of Neurology and the American Epilepsy Society. Neurology 84, 1705–1713 (2015).

11. Liu, G., Slater, N. & Perkins, A. Epilepsy: Treatment Options. Am Fam Physician 96, 87–96 (2017).

12. van Vliet, E.A., Aronica, E. & Gorter, J.A. Role of blood-brain barrier in temporal lobe epilepsy and pharmacoresistance. Neuroscience 277, 455–473 (2014).

13. Rodin, E.A., Rhodes, R.J. & Velarde, N.N. The prognosis for patients with epilepsy. J Occup Med 7, 560–563 (1965).

14. Lindsay, J., Ounsted, C. & Richards, P. Long-term outcome in children with temporal lobe seizures. I: Social outcome and childhood factors. Dev Med Child Neurol 21, 285–298 (1979).

15. Lindsay, J., Ounsted, C. & Richards, P. Long-term outcome in children with temporal lobe seizures. II: Marriage, parenthood and sexual indifference. Dev Med Child Neurol 21, 433–440 (1979).

16. Engel, J., Jr. What can we do for people with drug-resistant epilepsy? The 2016 Wartenberg Lecture. Neurology 87, 2483–2489 (2016).

17. Villeneuve, N. [Quality-of-life scales for patients with drug-resistant partial epilepsy]. Rev Neurol (Paris) 160 Spec No 1, 5S376–393 (2004).

18. DeLong, M. & Wichmann, T. Deep brain stimulation for movement and other neurologic disorders. Ann N Y Acad Sci 1265, 1–8 (2012).

19. Anderson, W.S. & Lenz, F.A. Surgery insight: Deep brain stimulation for movement disorders. Nat Clin Pract Neurol 2, 310–320 (2006).

20. Hosobuchi, Y., Adams, J.E. & Rutkin, B. Chronic thalamic and internal capsule stimulation for the control of central pain. Surg Neurol 4, 91–92 (1975).

21. Velasco, M., et al. Subacute electrical stimulation of the hippocampus blocks intractable temporal lobe seizures and paroxysmal EEG activities. Epilepsia 41, 158–169 (2000).

22. Kim, H.K., et al. Optogenetic intervention of seizures improves spatial memory in a mouse model of chronic temporal lobe epilepsy. Epilepsia 61, 561–571 (2020).

23. Stefan, H., et al. Magnetic source localization in focal epilepsy. Multichannel magnetoencephalography correlated with magnetic resonance brain imaging. Brain 113 (Pt 5), 1347–1359 (1990).

24. Fischer, M.J., Scheler, G. & Stefan, H. Utilization of magnetoencephalography results to obtain favourable outcomes in epilepsy surgery. Brain 128, 153–157 (2005).

25. Englot, D.J., et al. Epileptogenic zone localization using magnetoencephalography predicts seizure freedom in epilepsy surgery. Epilepsia 56, 949–958 (2015).

26. Baumgartner, C., Pataraia, E., Lindinger, G. & Deecke, L. Magnetoencephalography in focal epilepsy. Epilepsia 41 Suppl 3, S39–47 (2000).

27. Ben-Ari, Y., Tremblay, E. & Ottersen, O.P. Injections of kainic acid into the amygdaloid complex of the rat: an electrographic, clinical and histological study in relation to the pathology of epilepsy. Neuroscience 5, 515–528 (1980).

28. Krishnan, V., et al. Wireless control of cellular function by activation of a novel protein responsive to electromagnetic fields. Sci Rep 8, 8764 (2018).

29. Hunt, R.D., et al. Swimming direction of the glass catfish is responsive to magnetic stimulation. PLoS One 16, e0248141 (2021).

30. Cywiak, C., et al. Non-invasive neuromodulation using rTMS and the electromagnetic-perceptive gene (EPG) facilitates plasticity after nerve injury. Brain Stimul 13, 1774–1783 (2020).

31. Ashbaugh, R.C., Udpa, L., Israeli, R.R., Gilad, A.A. & Pelled, G. Bioelectromagnetic Platform for Cell, Tissue, and In Vivo Stimulation. Biosensors (Basel) 11(2021).

32. Hwang, J., et al. Regulation of Electromagnetic Perceptive Gene Using Ferromagnetic Particles for the External Control of Calcium Ion Transport. Biomolecules 10(2020).

33. Dimidschstein, J., et al. A viral strategy for targeting and manipulating interneurons across vertebrate species. Nat Neurosci 19, 1743–1749 (2016).

34. Barth, D.S., Sutherling, W., Engle, J., Jr. & Beatty, J. Neuromagnetic evidence of spatially distributed sources underlying epileptiform spikes in the human brain. Science 223, 293–296 (1984).

35. Sutherling, W.W., et al. The magnetic field of complex partial seizures agrees with intracranial localizations. Ann Neurol 21, 548–558 (1987).

36. Eliashiv, D.S., Elsas, S.M., Squires, K., Fried, I. & Engel, J., Jr. Ictal magnetic source imaging as a localizing tool in partial epilepsy. Neurology 59, 1600–1610 (2002).

37. Hari, R. & Kaukoranta, E. Neuromagnetic studies of somatosensory system: principles and examples. Prog Neurobiol 24, 233–256 (1985).

38. Knowlton, R.C. & Shih, J. Magnetoencephalography in epilepsy. Epilepsia 45 Suppl 4, 61–71 (2004).

39. Chabardes, S., et al. Deep brain stimulation in epilepsy with particular reference to the subthalamic nucleus. Epileptic Disord 4 Suppl 3, S83–93 (2002).

40. Jin, H., et al. Hippocampal deep brain stimulation in nonlesional refractory mesial temporal lobe epilepsy. Seizure 37, 1–7 (2016).

41. Usui, N., et al. Suppression of secondary generalization of limbic seizures by stimulation of subthalamic nucleus in rats. J Neurosurg 102, 1122–1129 (2005).

42. Yu, T., et al. High-frequency stimulation of anterior nucleus of thalamus desynchronizes epileptic network in humans. Brain 141, 2631–2643 (2018).

43. Morrell, M. Brain stimulation for epilepsy: can scheduled or responsive neurostimulation stop seizures? Curr Opin Neurol 19, 164–168 (2006).

44. Skarpaas, T.L., Jarosiewicz, B. & Morrell, M.J. Brain-responsive neurostimulation for epilepsy (RNS((R)) System). Epilepsy Res 153, 68–70 (2019).

45. Jarosiewicz, B. & Morrell, M. The RNS System: brain-responsive neurostimulation for the treatment of epilepsy. Expert Rev Med Devices 18, 129–138 (2021).

46. Schultz, D.M., et al. Sensor-driven position-adaptive spinal cord stimulation for chronic pain. Pain Physician 15, 1–12 (2012).

47. Pelkey, K.A., et al. Hippocampal GABAergic Inhibitory Interneurons. Physiol Rev 97, 1619–1747 (2017).

48. Haberman, R., et al. Therapeutic liabilities of in vivo viral vector tropism: adeno-associated virus vectors, NMDAR1 antisense, and focal seizure sensitivity. Mol Ther 6, 495–500 (2002).

49. Veliskova, J., Velsek, L. & Moshe, S.L. Subthalamic nucleus: a new anticonvulsant site in the brain. Neuroreport 7, 1786–1788 (1996).

50. Deransart, C., Le, B.T., Marescaux, C. & Depaulis, A. Role of the subthalamo-nigral input in the control of amygdala-kindled seizures in the rat. Brain Res 807, 78–83 (1998).

51. Dybdal, D. & Gale, K. Postural and anticonvulsant effects of inhibition of the rat subthalamic nucleus. J Neurosci 20, 6728–6733 (2000).

52. Smith, E.J., et al. Use of high-content imaging to quantify transduction of AAV-PHP viruses in the brain following systemic delivery. Brain Commun 3, fcab105 (2021).

53. Paxinos, G., Watson, C.R. & Emson, P.C. AChE-stained horizontal sections of the rat brain in stereotaxic coordinates. J Neurosci Methods 3, 129–149 (1980).

54. Li, N., van Zijl, P., Thakor, N. & Pelled, G. Study of the Spatial Correlation Between Neuronal Activity and BOLD fMRI Responses Evoked by Sensory and Channelrhodopsin-2 Stimulation in the Rat Somatosensory Cortex. Journal of Molecular Neuroscience 53, 553–561 (2014).

55. Zhong, M., et al. Multi-session delivery of synchronous rTMS and sensory stimulation induces long-term plasticity. Brain Stimul 14, 884–894 (2021).

56. Jouroukhin, Y., Nonyane, B.A.S., Gilad, A.A. & Pelled, G. Molecular Neuroimaging of Post-Injury Plasticity. Journal of Molecular Neuroscience 54, 630–638 (2014).

57. Krishnan, V.S., et al. Multimodal Evaluation of TMS - Induced Somatosensory Plasticity and Behavioral Recovery in Rats With Contusion Spinal Cord Injury. Front Neurosci 13, 387 (2019).

58. Li, N., et al. Optogenetic-guided cortical plasticity after nerve injury. Proc. Natl. Acad. Sci. U.S.A. 108, 8838–8843 (2011).

59. Lu, H., et al. Transcranial magnetic stimulation facilitates neurorehabilitation after pediatric traumatic brain injury. Scientific Reports 5(2015).

